# GeNet: Deep Representations for Metagenomics

**DOI:** 10.1101/537795

**Authors:** Mateo Rojas-Carulla, Ilya Tolstikhin, Guillermo Luque, Nicholas Youngblut, Ruth Ley, Bernhard Schölkopf

## Abstract

We introduce GeNet, a method for shotgun metagenomic classification from raw DNA sequences that exploits the known hierarchical structure between labels for training. We provide a comparison with state-of-the-art methods Kraken and Centrifuge on datasets obtained from several sequencing technologies, in which dataset shift occurs. We show that GeNet obtains competitive precision and good recall, with orders of magnitude less memory requirements. Moreover, we show that a linear model trained on top of representations learned by GeNet achieves recall comparable to state-of-the-art methods on the aforementioned datasets, and achieves over 90% accuracy in a challenging pathogen detection problem. This provides evidence of the usefulness of the representations learned by GeNet for downstream biological tasks.

## 1. Introduction

The last two decades have seen an exponential decrease in the cost of next generation DNA sequencing, transforming the field of microbiome science (Turnbaugh et al., 2007; Pasolli et al., 2019). A *microbiota* is a community of microorganisms residing in a multi-cellular organism, and the *microbiome* is the collective genetic material of this microbiota. Recently, mechanisms by which the human microbiota has an effect on a variety of health outcomes have been discovered (Sonnenburg & Bäckhed, 2016), and its responsiveness to dietary and lifestyle interventions (Walker et al., 2011; Wu et al., 2011) may lead to effective ways to prevent disease and improve health outcomes. Given a biological sample, studying the effect of the microbiota on its host requires as a first step understanding *which* microorganisms it contains. Nonetheless, the output of sequencing technologies are DNA *reads*, noisy substrings of the genomes present in the biological sample. How does one process such a collection of reads to understand which organisms are present, and in which amounts? The problem of *shotgun metagenomic classification* aims to assign to each read the corresponding host organism, and is an essential first step before downstream analysis can be carried-out. Currently, state-of-the-art methods such as Centrifuge (Kim et al., 2016) and Kraken (Wood & Salzberg, 2014) rely on sequence alignment, and match each read against a large database of known genomes. This requires high amounts of memory for storing such databases, and becomes challenging as the amount of noise in the reads increases.

The availability of more affordable sequencing technologies has been accompanied by noisier reads. Oxford Nanopore’s MinION (Jain et al., 2016) comes with error rates close to 10%, orders of magnitude higher than typical noise levels for Illumina, a more expensive technology. Sequence alignment based methods suffer from increasing ambiguity as noise increases. Machine learning systems, on the other hand, can *learn* from the noise distribution of the input reads. Moreover, a classification model learns a *mapping* from input read to class probabilities, and thus does not require a database at run-time. Finally, machine learning systems provide *representations* of DNA sequences which can be leveraged for downstream tasks.

### Contributions

In this paper, we introduce GeNet, a convolutional neural network model for shotgun metagenomic classification. GeNet is trained end-to-end from *raw* DNA sequences and exploits a hierarchical taxonomy between organisms using a novel architecture. We compare GeNet against state-of-the-art methods on real datasets, and show that GeNet achieves similar precision and good recall, despite the occurrence of strong dataset shift. We show in two examples that the representations of DNA sequences learned by GeNet can be successfully exploited by a linear classifier, achieving over 90% recall at the species level in the introduced datasets and over 90% accuracy in a challenging pathogen detection problem, outperforming baseline features. Finally, the trained weights only require 126MB of memory which is orders of magnitude smaller than databases for current metagenomic classifiers, providing advantages for portable, affordable technologies targeted for field use.

## 2. Problem statement and model

### 2.1. Metagenomic classification

We consider the problem of *shotgun metagenomic classification* from raw DNA sequences. We are given a collection of organisms *G*_1_*, …, G*_*k*_ for which the complete genome is known^1^. Each genome is a string consisting of four *nucleotides* A, C, T and G. Moreover, a hierarchical taxonomy 𝒯 encoding the similarities between the *K* genomes is available. 𝒯 is a tree, and each of the *K* genomes lies in one of its leaves. There are *L* levels in the tree corresponding to different *taxonomic ranks*, ranking from coarser (e.g., *Life*) to finer (e.g., *Species*). Each level *𝓁* contains *N*_*𝓁*_ nodes, also called *taxa* (*taxon* in singular). The taxonomic ranks used for training GeNet and the corresponding number of taxa are depicted in Figure 1, and represent only a fraction of the known biological taxonomy. As an illustration, humans and chimpanzees are Homonidae, and thus both belong to the same taxonomic rank *Family*. Nonetheless, they both belong to a different *Genus*, and thus to a different *Species*, two finer taxonomic ranks.

**Figure 1.**
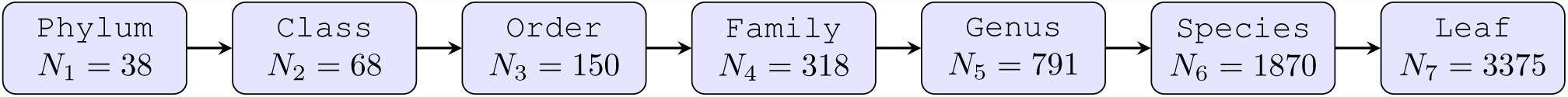
Taxonomic ranks in the tree 𝒯 used to train GeNet, and the number of taxa (nodes) at each level. The coarser level considered is *Phylum*, and the finer level before the leaves is *Species*. A taxonomic tree containing all living organisms has significantly more taxa.

When sequencing DNA from a biological sample, a specific *sequencing technology* is used. Some widely used technologies include Illumina, Pacbio and Nanopore. The output of these technologies are called *reads* and are noisy substrings **s** of the genomes present in the sample. The length of the reads and the noise they contain is characteristic to the sequencing technology, and results in nucleotides being flipped, deleted or added. The distribution of this noise, which we call *technology specific noise*, is in practice empirically estimated (McElroy et al., 2012). Illumina technologies produce short reads (around 100 nucleotides), all of the same length, while Nanopore technologies produce longer reads of varying length (roughly between 1, 000 and 10, 000 nucleotides).

Given an input read **s** which is a noisy substring of genome *G*_*k*_, 𝒯 defines a unique labeling **y** = (*y*_1_*, …, y*_*L*_), where *y*_*𝓁*_ 1*, …, N*_*𝓁*_ is the correct label, or taxon, at the *𝓁*-th level of 𝒯. The goal of **metagenomic classification** is to predict the correct taxon for a read **s** at different levels in 𝒯. For the example in Figure 1, metagenomic classification at the Phylum level consists of finding the correct taxon for read **s** among *N*_1_ = 38 possibilities.

If one is able to correctly label a read to the corresponding host genome *G*_*k*_, the correct hierarchical labelling vector **y** can be directly deduced by tracing the unique path to the root in 𝒯. However, it may not be possible to correctly classify a sequence at the finer levels of the tree. Genomes close in the taxonomy may share significant portions of their DNA, and reads from such genomes lead to ambiguous classification, especially at the finer levels of the taxonomic tree (Jain et al., 2018). Classification of short reads is also often ambiguous, and is still desirable to classify at coarser levels in the tree. Of particular interest when analysing biological properties of a sample are the levels of *genus* and *species*. For example, the study of pathogenesis (the biological mechanisms leading to disease) requires species level classification.

### 2.2. GeNet: a convolutional model for metagenomic classification

We propose GeNet, a model for metagenomic classification based on convolutional neural networks. GeNet is trained end-to-end from raw DNA sequences using standard backpropagation with cross-entropy loss. Given an input sequence **s** *∈* {0, 1, 2, 3}^*d*^, we first extract features **h** = *G*(**s**) *∈* ℝ^*q*^ using a Resnet like model (He et al., 2016) and a fully connected layer. This representation **h** is then mapped to *L* softmax layers, each of size *N*_*𝓁*_, which provides a probability distribution over the *N*_*𝓁*_ possible taxa at each level *𝓁* of the taxonomy 𝒯. The number of known genomes is significantly higher than could be easily handled by standard neural network architectures, since it would result in hundreds of thousands of classes. We train GeNet with a representative subset of all the known genomes, details are given in Section 4.

Denote by 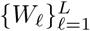 the weight matrices mapping **h** to softmax vectors of size *N*_*𝓁*_, the number of possible taxa at the *𝓁*-th level of the tree, where 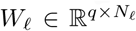. We allow for predictions at level *𝓁 -* 1 to inform predictions at level *𝓁*. For example, knowing that a read is likely to come from a bacterial organism narrows down the possible taxa at finer levels in 𝒯. To that end, we include in our model matrices 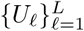 which allow that the unnormalised probability vector over taxa at level *𝓁 -* 1 contributes to the unnormalised probability vector at level *𝓁*, where 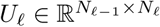 and *U*_0_ is a zero matrix. This leads to the following unnormalised probability vector over labels at each level of 𝒯 :

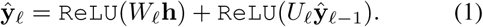

A diagram of GeNet is provided in Appendix A.

#### Training pipeline

GeNet receives as input a noise parameter *p*, or *base-calling probability*, and the size *r*_*max*_ of input reads. We choose to train with uniform noise, that is, each nucleotide in a read is flipped with probability *p*, and is replaced uniformly among the remaining three nucleotides. This allows us to remain agnostic to technology specific noise, and can be partially corrected after training if necessary as described in Section 2.3. During training, the genomes *G*_1_*, …, G*_*K*_ are stored in memory and new mini-batches are generated on-the-fly. For each iteration of stochastic gradient descent, we produce a new mini-batch of size *M* by randomly selecting *M* genomes among the *K* available, selecting a random location in each and extracting a read of varying length from this location. We add uniform noise with parameter *p* to each read, and pad it with zeroes to fit into the fixed input length *r*_*max*_. For reads longer than the input length, only the first *r*_*max*_ letters of longer reads are considered^2^. This allows us to classify reads of varying length. Uniform noise was chosen to remain agnostic to technology specific noise. The procedure for sampling a mini-batch is described in Algorithm 1.

#### Loss function

Let 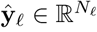 be the unnormalised probability vector predicted by the network over the taxa at level *𝓁* as defined in Equation 1, and let **y** = (*y*_1_*, …, y*_*L*_) be the true taxa for the input sequence **s** at each level of the taxonomy. We denote by **y**_*𝓁*_ the one-hot encoding of *y*_*𝓁*_, so that **y**_*𝓁*_ is a zero vector of size *N*_*𝓁*_ except for location *y*_*𝓁*_, which equals one.

At each level of 𝒯, we compute the cross-entropy loss

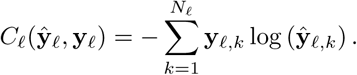

The distribution of taxa in 𝒯 gives rise to class imbalance which we found necessary to correct for successful training. To illustrate, assume that *K* genomes are seen uniformly during training. Consider now a coarser level in the tree, such as *Domain* (not used to train GeNet), which contains the taxa *Bacteria*, *Virus* and *Archaea*. For this illustration, assume that of the *K* genomes, 90% are bacteria, 8% virus and 2% archaea, meaning that the taxa at the *Domain* level are highly imbalanced. Similar imbalances are likely to appear at most levels of the tree. If this imbalance is not corrected, any non bacterial organism wrongly classified as a bacteria is likely to be wrongly classified at finer levels of 𝒯. Therefore, each level of the tree uses a vector 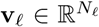 which down-weights the contribution to the loss of more abundant taxa. In the previous example, **v**_*Domain*_ = (1*/*0.9, 1*/*0.08, 1*/*0.02). If a uniform distribution is assumed at the leaf level, the resulting *L* weight vectors are determined by the *K* genomes and the tree 𝒯.

The overall loss is then the weighted sum of cross-entropy losses at all the levels in the tree,

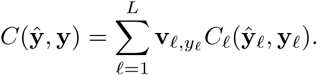

GeNet is trained using standard stochastic gradient descent with Nesterov momentum (Sutskever et al., 2013).

#### Architecture choice

Recurrent neural networks are the standard tool when considering sequential inputs. Nonetheless, we chose a convolutional architecture mainly due to i) speed constraints and ii) more success achieving high validation accuracies. We found that achieving good coverage of the *K* genomes is important to achieve high validation accuracies, i.e., we must observe most parts of all the genomes. The combined length of the genomes used to train GeNet is over 10 billion and GeNet trained for about one week on a NVIDIA P-100 GPU until convergence. The input to the network is a sum of a one-hot encoding of the letters in the sequence, an embedding of the sequence and a positional embedding as proposed in Gehring et al. (2017). Both the embedding matrix and positional embedding matrix are learned during training. Many applications involving sequential data have benefited from convolutional architectures, for example machine translation (Gehring et al., 2017), text classification (Conneau et al., 2017) and video classification (Karpathy et al., 2014).

### 2.3. Domain adaptation and general purpose representations

Since any test dataset obtained from real sequencing technologies represents a probability distribution that differs from the training distribution of GeNet, we face dataset shift. First, the noise distribution of real reads is not uniform, and depends on the sequencing technology. Moreover, the proportion of genomes is often not uniform, since longer genomes are over-represented, and the sample contains genomes in different proportions altogether. Genomes which were not observed during training may also be present. A priori, there are therefore no guarantees regarding the generalization performance of GeNet to unseen data if no further training takes place. Experiments in Section 4.1 report generalization performance of GeNet on real datasets.

#### Algorithm 1 SAMPLE MINI-BATCH

**Input:** Reference genomes *G*_1_*, …, G*_*K*_. Taxonomic tree 𝒯. Input size *r*_*max*_, minimum read length *r*_*min*_, base-calling error probability *p*. Mini-batch size *M*.

**Returns:** Mini-batch of noisy sequences *S* and corresponding hierarchical labels *Y.*

Initialize *S* =[ ], *Y* =[ ].

**for** *j* = 1 **to** *M* **do**

   Select a *genome G*_*k*_ uniformly at random.

   Select a *location j ∈*{1*, …, L*_*k*_ *- r*_*max*_} uniformly at random, where *L*_*k*_ is the length of *G*_*k*_.

   Select a *read length r ∈*{*r*_*min*_*, …, r*_*max*_} uniformly at random.

   Define the read **s**_*j*_ = *G*_*k*_(*i* : *i* + *r*), add uniform flipping noise with probability *p* and pad with zeroes at the end of the read, so that **s**_*j*_ has length *r*_*max*_.

   Obtain the corresponding *label vector* **y**_*j*_ from 𝒯.

   *S.*add(**s**_*j*_), *Y.*add(**y**_*j*_)

**end for**

Nonetheless, intermediate activations of the network provide representations of the DNA reads which can be used for downstream tasks. Given a supervised learning problem with training data 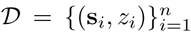, where **s** are DNA sequences and *z* are outcomes we wish to predict from these sequences, we propose to use GeNet to compute representations of the training inputs **h**_*i*_ = *G*(**s**_*i*_), where **h**_*i*_ is the last hidden layer of GeNet, see Section 2.2. We then train a classification model on the transformed dataset 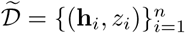 and evaluate test performance.

We show that the representations in the last hidden layer of GeNet can be used for downstream tasks in Section 4.2 in two different examples. First, we consider datasets of Nanopore reads, for which dataset shift occurs, and we show that a linear model trained on GeNet representations achieves classification accuracy competitive with the state-of-the-art. Second, we consider the binary classification problem of deciding whether a read comes from a pathogenic organism, several of which were not observed during training, and show that a linear model achieves over 90% test accuracy. Such use of pre-trained features has had remarkable success in image recognition (see Rawat & Wang (2017) and references therein) and natural language processing with representations such as word2vec (Mikolov et al., 2013). It is reassuring that the same holds true for metagenomics.

## 3. Related work

State-of-the-art methods for metagenomic classification rely on sequence alignment. They use a large database of known genomes *G*_1_*, …, G*_*K*_ and given a read **s**, do an exhaustive search to find one or more genomes *G*_*k*_ such that **s** is a substring of *G*_*k*_. Since reads are often noisy, and the genomes in the sample may not exactly match any genome in the reference database, a notion of distance is used to compare strings. BLAST (Altschul et al., 1990) performs this exhaustive search, but does not scale to real world datasets with millions of reads. State-of-the-art methods compress the database of genomes and strike a balance between accuracy and speed. These methods include Kraken (Wood & Salzberg, 2014) and Centrifuge (Kim et al., 2016).

The machine learning perspective on metagenomic classification is not new. Busia et al. (2018) build a convolutional network for metagenomic classification of short reads (under 200 nucleotides) on 16S data, and achieve good performance on a series of datasets. These data come from very specific parts of a conserved gene only present in prokaryotic organisms, excluding viruses and eukaryotes. Our approach focuses on the shotgun setting and supports reads originating from any part of the genome. Moreover, Busia et al. (2018) do not analyse the re-usability of features learned by the network for downstream tasks. Nissen et al. (2018) introduce a method for the related problem of metagenomic binning using Variational Autoencoeders (VAE) (Kingma & Welling, 2013), and show that the representations learned by the VAE are useful for clustering. Their system is trained on co-abundance and composition data, not raw DNA sequences. To our knowledge, GeNet is the first system trained from raw DNA sequences for shotgun metagenomics. Finally, Feng et al. (2018) introduce a deep learning system exploiting function hierarchy for gene function prediction.

In machine learning, using label hierarchy to boost classification performance is not new. Silla & Freitas (2011) provide a survey of standard methods used by practitioners in different fields. One option is to flatten the tree, essentially performing classification at the leaf node and disregarding the taxonomy. The availability of the tree does however allow for hierarchical classification during evaluation, since a correctly classified leaf is correct at all levels in the taxonomy. Another approach is having local classifiers at different nodes in the tree (Vural & Dy, 2004; Cerri et al., 2014), often of limited applicability for hierarchies with a large number of nodes. In metagenomic classification, such an approach would suffer from short read lengths for which ambiguity is high, since the likelihood of a sequence belonging to several unrelated genomes becomes higher. Babbar et al. (2013) provide data-dependent bounds indicating when exploiting the hierarchy helps compared to using the flattened tree. Levatić et al. (2015) also analyse the usefulness of exploiting label hierarchy in a wide range of classification tasks. Recent papers have encoded the hierarchy in a neural network architecture. Zhu & Bain (2017) propose a multi-branch convolutional network, in which each branch aims to predict the label at a specific level of the tree similarly to GeNet. However, contrary to this approach, GeNet predicts labels at all levels of the tree from the same hidden representation simultaneously.

## 4. Experiments

We provide experimental analysis of GeNet in two different problems. Section 4.1 analyses the generalization ability of GeNet to reads for which dataset shift occurs. Section 4.2 shows that the representations learned by GeNet are useful for downstream tasks, a fundamental advantage.

We compare GeNet to two state-of-the-art methods for metagenomic classification: Centrifuge (Kim et al., 2016) and Kraken2 (Wood & Salzberg, 2014)^3^. We now introduce the training dataset and training details. The code to train and evaluate GeNet will be made available at https://github.com/mrojascarulla/GeNet.

### Training dataset and parameters

We use a variation of the dataset from Kim et al. (2016) to train GeNet as described in Section 2.2. This dataset consists of 4278 prokaryotic genomes available in RefSeq (Pruitt et al., 2013). To avoid redundancies, we removed genomes which shared the same taxonomy at the leaf level. This resulted in a dataset with *K* = 3375 genomes, whose NCBI reference IDs may be found at https://github.com/mrojascarulla/GeNet.

These *K* genomes are loaded in memory and GeNet is trained, with mini-batches generated as described in Algorithm 1. We trained i) three networks with different values of the noise parameter *p ∈*{0.03, 0.1, 0.2} for reads of fixed length *r*_*max*_ = *r*_*min*_ = 1000 and ii) a network with noise *p* = 0.1, *r*_*min*_ = 1000 and *r*_*max*_ = 10000. For each network, we carry out hyper-parameter search for some network parameters using validation data generated identically to the training data, see Table 8 in Appendix A.

We optimize the network with mini-batches of size 64 using Nesterov momentum (Sutskever et al., 2013), with momentum parameter 0.9. Training to convergence takes roughly one week on a P-100 GPU. Details on the architecture and hyper-parameters are provided in Appendix A.

### Performance metrics

We report three performance metrics. First, we consider recall. For a dataset with *p* classes, denote by *r*_*i*_ the one-vs-all recall measured for class *i* and *w*_*i*_ the proportion of examples corresponding to class *i* in the dataset. Then the overall recall is 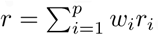. This is equivalent to standard accuracy. Similarly, we report precision 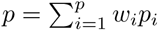, where *p*_*i*_ is the one-vs-all precision of class *i*.

We also report the Average Nucleotide Identity (ANI), computed using FastANI (Jain et al., 2018). ANI is a measure of similarity between two genomes. We report the ANI between the genome predicted by GeNet and the true genome, indicating if the network is making “sensible” mistakes by predicting incorrect but similar labels. FastANI returns zero for similarity values smaller than a given threshold. In such cases, we use Average Amino-Acid Identity (AAI) instead, computed with CompareM (Parks). Details for ANI and AAI computations are given in Appendix B.3.

Results are given both at the biological ranks (levels in the tree 𝒯) of *genus* and *species*, two of the finer levels of the taxonomy, see Figure 1. In addition to GeNet, we report GeNet top 5, in which we consider a read to be correctly classified if the true label is among the top 5 highest ranked predictions for that read.

### 4.1. Generalization to other data distributions

We first evaluate the performance of GeNet on data drawn from different distributions, on which domain shift occurs. We consider two problems of increasing difficulty.

First, we build a dataset with reads belonging to the *K* genomes used during training. These reads are generated with added Illumina-type noise using the random-reads module of BBMap (Bushnell), an open-source short read aligner. For this experiment, the domain shift only occurs in the distribution of the noise, not the distribution of the labels. We generate reads with Illumina noise levels *q ∈* {0.01, 0.1, 0.2, 0.3}. For each noise level, we generate ten datasets with a different random seed, each with one million reads split uniformly between the *K* genomes. We test with GeNet trained on uniform noise parameters *p* = 0.03 for *q* = 0.01, *p* = 0.1 for *q* = 0.1 and *p* = 0.2 for higher *q*. Figure 2 reports *per class* recall at the genus level. Conclusions from the recall at species level are similar. Details on the parameters of BBMap are given in Appendix B.1.

**Figure 2.**
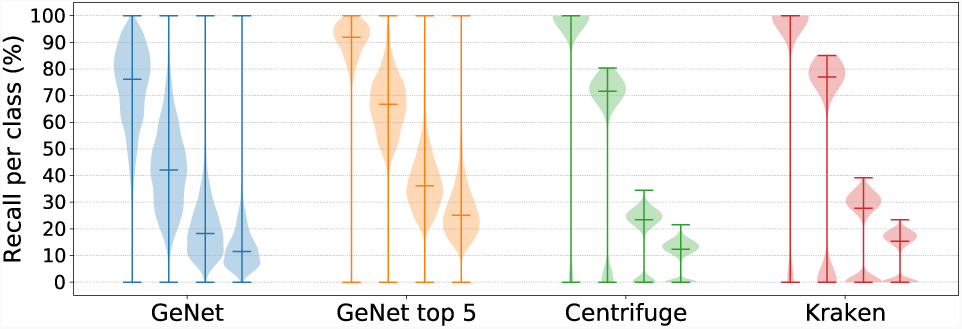
**Recall per class at the genus level on Illumina datasets.** For each method, four results are available, corresponding to datasets with increasing read noise *q ∈*{0.01, 0.1, 0.2, 0.3}. The median recall per class is reported. The average performance of Kraken and Centrifuge is higher than GeNet. For the two higher noise settings, GeNet top 5 outperforms Kraken and Centrifuge. As noise increases, the distribution of per class recall is bi-modal for Centrifuge and Kraken, and close to half of the labels have recall near zero, which is undesirable. The distribution of per class recall for GeNet is uni-modal, and remains similar as noise increases.

Second, we consider four real world datasets of Nanopore reads introduced in Nicholls et al. (2018), with the following accession numbers in the European Nucleotide Archive: `ERR2906227`, `ERR2906228`, `ERR2906229` and `ERR2906230`. Below, we call these datasets 𝒟1*, …,* 𝒟_4_. Both 𝒟_1_ and 𝒟_3_ contain around 3 million reads, while 𝒟_2_ and 𝒟_4_ contain over 30 million reads. These datasets contain reads from 10 organisms, of which we only consider 8 (the remaining two eukaryotes, Cryptococcus neoformans and Saccharomyces cerevisiae, were not observed during training, so GeNet cannot classify them into the correct class). To obtain ground truth labels, we map the reads to the genomes present in the dataset using minimap2 (Li, 2018), a DNA aligner. Details on the use of minimap2 are given in Appendix B.2. We report precision and recall in Tables 1 and 2 respectively. Moreover, we report Average Nucleotide Identity (ANI) between the predictions of GeNet and the true labels in Figure 4. We also analyse the effect of read length in accuracy in Figure 3.

**Table 1:** **Precision on the Nanopore datasets** from Nicholls et al. (2018) at two of the finer levels in the taxonomy 𝒯, genus and species. GeNet achieves high precision and is competitive with Kraken and Centrifuge.

| Method | $\mathcal{D}_1$ (%) | | $\mathcal{D}_2$ (%) | | $\mathcal{D}_3$ (%) | | $\mathcal{D}_4$ (%) | |
| --- | --- | --- | --- | --- | --- | --- | --- | --- |
|  | Genus | Species | Genus | Species | Genus | Species | Genus | Species |
| GeNet | 94.6 | 96.1 | 93.9 | 95.5 | 98.5 | 98.8 | 98.5 | 98.8 |
| GeNet top 5 | 97.7 | 98.2 | 97.6 | 98.0 | 99.1 | 99.1 | 98.9 | 99.1 |
| Kraken | 96.6 | 96.7 | 96.5 | 96.6 | 98.4 | 98.5 | 98.4 | 98.5 |
| Centrifuge | 97.4 | 97.4 | 97.2 | 97.3 | 98.4 | 98.6 | 98.3 | 98.6 |

**Table 2:**
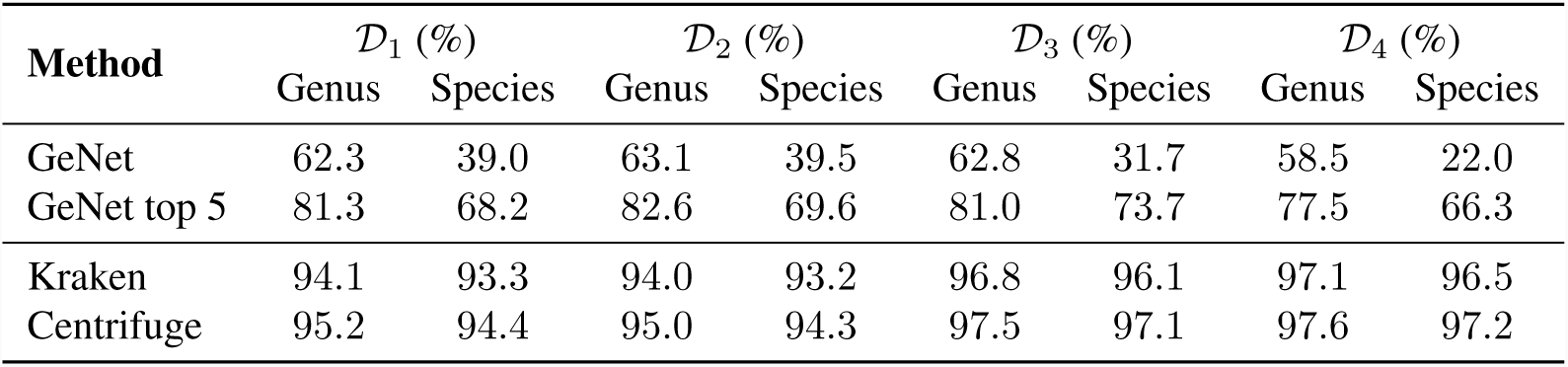
**Recall on the Nanopore datasets** from Nicholls et al. (2018) at the genus and species levels. Kraken and Centrifuge achieve significantly higher recall than GeNet, due to the strong dataset shift in the distribution of the labels and the noise distribution. For example, 93% of 𝒟_3_ is composed of one bacterial species only. GeNet still performs way above chance (0.13% at the genus level and 0.05% at the species level), testifying to a significant amount of domain adaptation.

**Figure 3.**
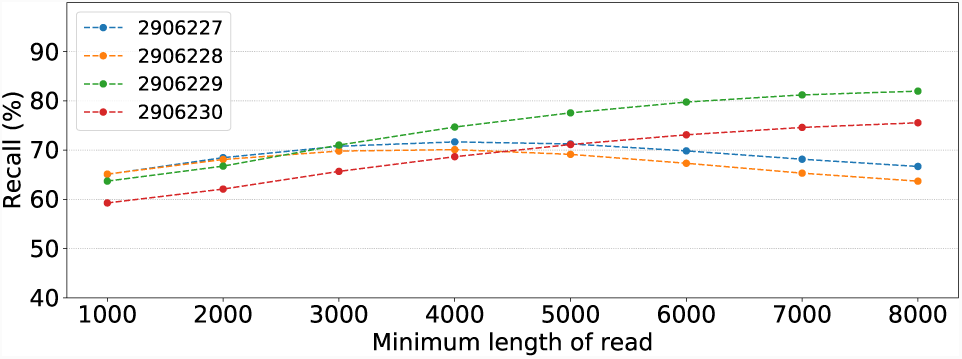
**Recall of GeNet at the genus level on the four Nanopore datasets** from Nicholls et al. (2018) as a function of minimal read length. For two datasets, GeNet performs better on longer reads, earning a 15% difference between all reads longer than 1, 000 to reads longer than 8, 000. Recall is roughly un-changed on the other two datasets.

**Figure 4.**
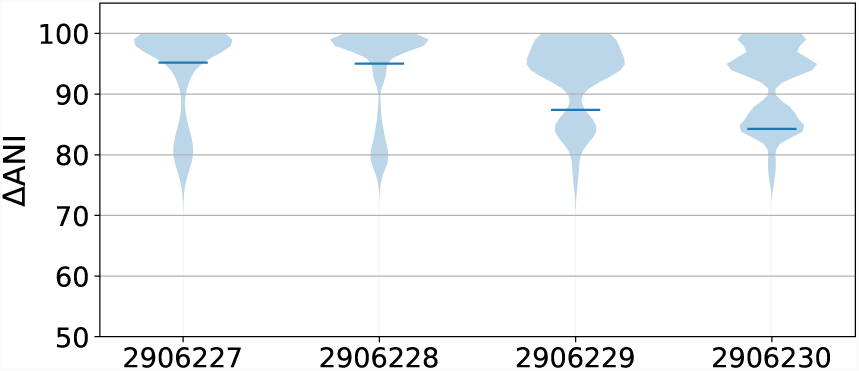
Average Nucleotide Identity (ANI) between genome the genome with the highest predicted probability for GeNet and the true genome on the four Nanopore datasets. For values of ANI smaller than 70, Amino Acid Identity (AAI) was used instead. Since GeNet was only trained with reads longer than 1, 000, we discard shorter reads in this plot, excluding roughly 7% of predictions in 𝒟_1_ and 𝒟_2_ and 1.5% on the other two datasets. For the first two datasets, median ANI is close to 95%, and closer to 85% for the other unbalanced datasets.

### 4.2. Downstream tasks with learned representations

We showcase the re-usability of the representations in the last hidden layer of GeNet in two downstream tasks as described in Section 2.3. Given a labelled dataset 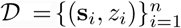, we train a linear logistic regression on the transformed dataset 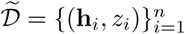, where the label *z*_*i*_ depends on the specific example and **h** *∈* ℝ^1024^. We denote this method *GeNet + LIN*. As an alternative representation of the reads, we divide a sequence **s**_*i*_ in ten equally sized bins, and compute the frequency of each of the four nucleotides in every bin. The resulting 40 dimensional vector 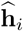 is an alternative representation of **s**_*i*_. As baselines, we train a logistic regression and a Multi Layer Perceptron on the transformed dataset 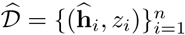 with grid search and cross validation, resulting in methods *Freq + LIN* and *Freq + MLP*.

As a first downstream task, we consider the four Nanopore datasets discussed in Section 4.1. We sub-sample each dataset to contain only 3 million reads for computational reasons. In each dataset separately, we use 10, 000 randomly chosen examples with their true labels *at the species level* as a labelled dataset 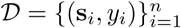. Note that this is only 0.33% of the size of each dataset. We evaluate performance on the remaining held-out reads and average the results over ten repetitions, see Table 4.

**Table 3:** **Accuracy in the pathogen detection problem.** 21 closely related organisms are considered, 11 of which are pathogenic. The logistic regression models and MLP are trained using a labeled set of size 10, 000, and accuracy is computed on the remaining held-out data. Results are averaged over 10 such repetitions. The recalls obtained by Kraken and Centrifuge on the whole dataset are also reported for reference. For pathogenesis, species level classification is necessary, so a prediction for these two methods is considered correct if the read is detected correctly at the species level. *GeNet + LIN* significantly outperform *Freq + LIN* and *Freq + MLP*, which exhibit near chance performance. While not directly comparable, since these methods do not have access to an additional training phase, *GeNet + LIN* outperforms Centrifuge on this task.

| Method | Accuracy |
| --- | --- |
| GeNet + LIN | **90.3 ± 0.1** |
| Freq + LIN | 50.8 ± 1.0 |
| Freq + MLP | 61.4 ± 1.4 |
| Centrifuge | 88.5 |
| Kraken | **99.5** |

**Table 4:**
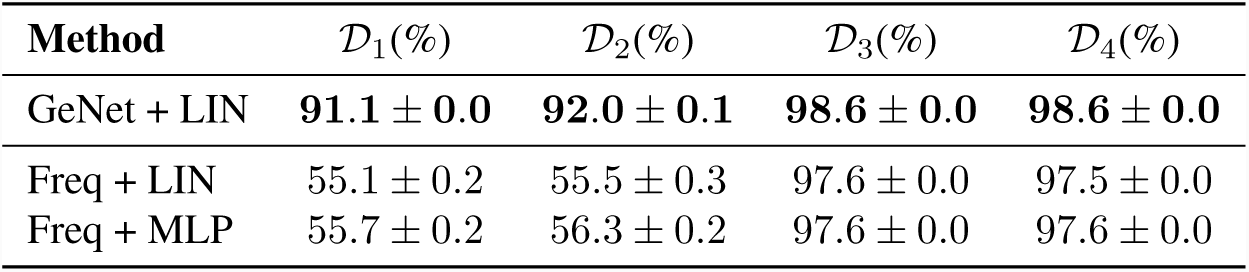
**Recall at the species level on the four Nanopore datasets.** GeNet is used to compute representations of the reads in the datasets, and a logistic regression is trained on a labeled dataset with 10, 000 reads. Features built using nucleotide frequencies are used as a benchmark, see Section 4.2. Recall is computed on the remaining held-out data. Averages over 10 repetitions are reported. *GeNet + LIN* significantly outperforms *Freq + LIN* and *Freq + MLP*, and is closer to the recalls obtained by Centrifuge and Kraken. *Freq + LIN* and *Freq + MLP* achieve high recall on two datasets, however, these datasets are highly unbalanced and are composed over 90% of one type of bacteria. These models always predict the majority class, and obtain zero recall for other classes. This is not the case for *GeNet + LIN*.

| Method | $\mathcal{D}_1(\%)$ | $\mathcal{D}_2(\%)$ | $\mathcal{D}_3(\%)$ | $\mathcal{D}_4(\%)$ |
| --- | --- | --- | --- | --- |
| GeNet + LIN | **91.1 ± 0.0** | **92.0 ± 0.1** | **98.6 ± 0.0** | **98.6 ± 0.0** |
| Freq + LIN | 55.1 ± 0.2 | 55.5 ± 0.3 | 97.6 ± 0.0 | 97.5 ± 0.0 |
| Freq + MLP | 55.7 ± 0.2 | 56.3 ± 0.2 | 97.6 ± 0.0 | 97.6 ± 0.0 |

As a second task, we build a dataset of 21 closely related genomes, of which 10 are highly pathogenic (harmful) and the rest are not known to cause any diseases. The dataset consists of 4 Clostridium strains, one Clostridiodes strain, 4 Vibrio strains, 5 Pseudomonas strains, 2 Klebsiella strains, 3 Streptococcus strains and 2 Burkholderia strains (each of these indicates a different genus). Within each genus, only some of the strains are pathogenic. The corresponding NCBI accession IDs can be found at https://github.com/mrojascarulla/GeNet. We consider the labelled dataset 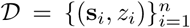 where *z*_*i*_ is a binary label indicating whether the corresponding read belongs to a pathogenic genome. The reads are generated using BBMap, with PacBio noise of 10%. The dataset has 100, 000 reads, of which 10, 000 randomly chosen reads are used for training. Accuracy is evaluated on the remaining held-out reads, averaged over 10 repetitions, see Table 3.

### 4.3. Analysis of the results

We draw the following conclusions from our experiments.

### GeNet partially generalises to real data

Our experiments show that despite the occurring dataset shift, GeNet can perform significantly above chance in real datasets at fine levels of the hierarchical taxonomy. For Illumina reads, the recall achieved by Kraken and Centrifuge is higher than that of GeNet. Nonetheless, Figure 2 shows that as noise increases, Kraken and Centrifuge achieve recall close to zero for a large proportion of the genomes. A more uniform recall distribution per genome may be desirable, and is partially achieved by GeNet for which few classes have recall close to zero, even as noise increases. This showcases the usefulness of training with noise.

On the Nanopore datasets, recall for GeNet is significantly lower than for Centrifuge and Kraken, see Table 2, while precision is competitive, see Table 1. We attribute the low recall mainly to the strong distribution shift, both in the output labels (only 8 genomes are present in the datasets, while we used 3375 for training, and 𝒟_3_ and 𝒟_4_ are highly unbalanced) and the noise distribution. Nonetheless, the ANI distribution in Figure 4 witnesses that often the predicted genomes are similar to the true genomes, which means that most mistakes are not unreasonable. Moreover, GeNet is expected to perform better as read length increases, which is the case in two of the datasets, see Figure 3.

### GeNet representations perform well in downstream tasks

Experiments on downstream tasks show that the representations learned by GeNet can be successfully exploited. While GeNet significantly under-performs state-ofthe-art methods in terms of recall on the Nanopore datasets, training a linear model on top of GeNet representations leads to a significant increase in performance in these datasets, see Table 4. While we cannot compare the results directly with Centrifuge and Kraken since these do not have access to an extra training phase, the obtained recalls are competitive. In practice, a small percentage of a target dataset of reads can be labeled with alternative methods to train such a supervised model.

Second, using standard frequency features on the pathogen dataset leads to close to chance performance. This showcases the difficulty of this problem from raw data. *GeNet + LIN* achieves over 90% held-out accuracy, see Table 3. While direct comparison with Kraken and Centrifuge is not possible in this case, it is encouraging that *GeNet + LIN* outperforms Centrifuge in this problem.

### GeNet strikes a trade-off between speed and storage

Computing predictions for reads of size 10, 000 on a NVIDIA P-100 is roughly 10 times slower than Centrifuge and 20 times slower than Kraken, see Table 5. Nonetheless, the only storage required by GeNet are the weights of the network, which require 126MB. This is two orders of magnitude smaller than Centrifuge, and three orders of magnitude smaller than Kraken.

**Table 5:** Speed and memory requirements of metagenomic classifiers. Speed is computed during the evaluation of dataset 𝒟_3_, which contains over 3 million reads. We report speed for GeNet using both P-40 and P-100 GPU cards. Kraken uses 8 CPU cores and 120 GB of RAM. Speed for Centrifuge is reported from Kim et al. (2016), since Centrifuge does not return how much time was spent loading the database and how much classifying the reads. GeNet is slower than the competitors, inference is only an order of magnitude slower than for Centrifuge. However, the memory usage of GeNet is smaller by two orders of magnitude.

| Method | Speed (reads/min) | Memory (GB) |
| --- | --- | --- |
| GeNet, P-100 | 71,000 | **0.126** |
| GeNet, P-40 | 57,000 | - |
| Kraken | **1,732,000** | 93.0 |
| Centrifuge | 563,380 | 11.3 |

## 5. Conclusion

We showed in two datasets obtained from different sequencing technologies that GeNet achieves good recall and high precision despite strong dataset shift. We also provided evidence that the representations in the last hidden layer of GeNet can be used for downstream tasks. Training a linear model with a small percentage of labelled data in Nanopore datasets leads to an increase of recall at the species level of 50% or more. Moreover, GeNet features significantly outperform frequency features computed from the raw sequences on a challenging pathogen detection problem.

We expect that GeNet representations can be used for a variety of tasks in computational biology, e.g., gene function prediction. Many tasks require a *representation* of the data and thus cannot benefit from methods such as Kraken and Centrifuge, notwithstanding the excellent performance these custom tools exhibit in metagenomic classification. In addition, our approach exhibits a higher level of noise robustness, the ability to learn from technology specific noise, as well as small memory requirements. This can present interesting opportunities for the development of cheaper and portable sequencing technologies.

The use of pre-trained networks and data representations has accelerated research in computer vision, speech and natural language processing, allowing fast deployment of solutions for new problems that often come with small labelled training sets. We anticipate a similar potential for computational biology and health, where labelled data sets can also he hard to come by. The ultimate promise and validation of the proposed method would thus consist of its adoption by the community and application in a diverse array of tasks, which is well beyond the scope of the present work.

## Acknowledgements

The authors are thankful to Sylvain Gelly for helpful discussions on the architecture design for GeNet.

## Appendix to “GeNet: Deep Representations for Metagenomics”

### A. Training details

GeNet was implemented using Tensorflow (Abadi et al., 2015).

The vocabulary size is 6: four nucleotides `A`,`C`, `T` and `G`, an end-of-sequence character, and a character for *ambiguous* nucleotides, which appears occasionally on the downloaded genomes. The input to GeNet is a sum of the one-hot encoding of each letter in the input sequence, a trainable six dimensional embedding, and a trainable six dimensional positional embedding as proposed in Gehring et al. (2017). For an input sequence of length *r*_*max*_, this results in a matrix of shape 6 *× r*_*max*_. The architecture of GeNet is available in Table 7 and is depicted in Figure 5. Details for the Resnet blocks used can be found in Table 6. For every call of a Resnet block of the form (*n,* 2*n*), the number of filters is multiplied by two, and the size of the input is divided by two. When there is a size mismatch between the input to the Resnet block and the output, a 1d convolution with the appropriate number of filters is used to match the dimensions.

**Table 6:** Resnet block used in GeNet. The input is added to CONV in the last layer using a 1 × 1 convolution with 2*n* filters. Resnet blocks of the form (*n, n*) do not perform average pooling as a first s tage, s o t he i nput a nd output number of filters is unchanged.

| Resnet block $n \times 2n$ |
| --- |
| Layers |
| AVG. POOL $1 \times 2$, stride 2.<br>BN + ReLU. |
| CONV $1 \times w \times 2n$, stride 1.<br>BN + ReLU. |
| CONV $1 \times w \times 2n$, stride 1.<br>Input + CONV |

**Table 7:** Layers of GeNet. `BN` stands for Batch Norm (Ioffe & Szegedy, 2015), `FC` for fully connected. `CONV` *v × w × n*_*f*_ stands for a 2d convolutional layer with kernel size (*v, w*) and *n*_*f*_ filters.

| GeNet |
| --- |
| Layers |
| Embedding + Pos.embedding + One-hot.<br>CONV $v \times w \times n_f$, stride $w$.<br>RESNET BLOCK $n_f \times n_f$<br>RESNET BLOCK $n_f \times n_f$<br>RESNET BLOCK $n_f \times 2n_f$<br>RESNET BLOCK $2n_f \times 2n_f$<br>BN + ReLU<br>AVG. POOL + BN<br>FC $f_c$.<br>L softmax layers. |

**Figure 5.**
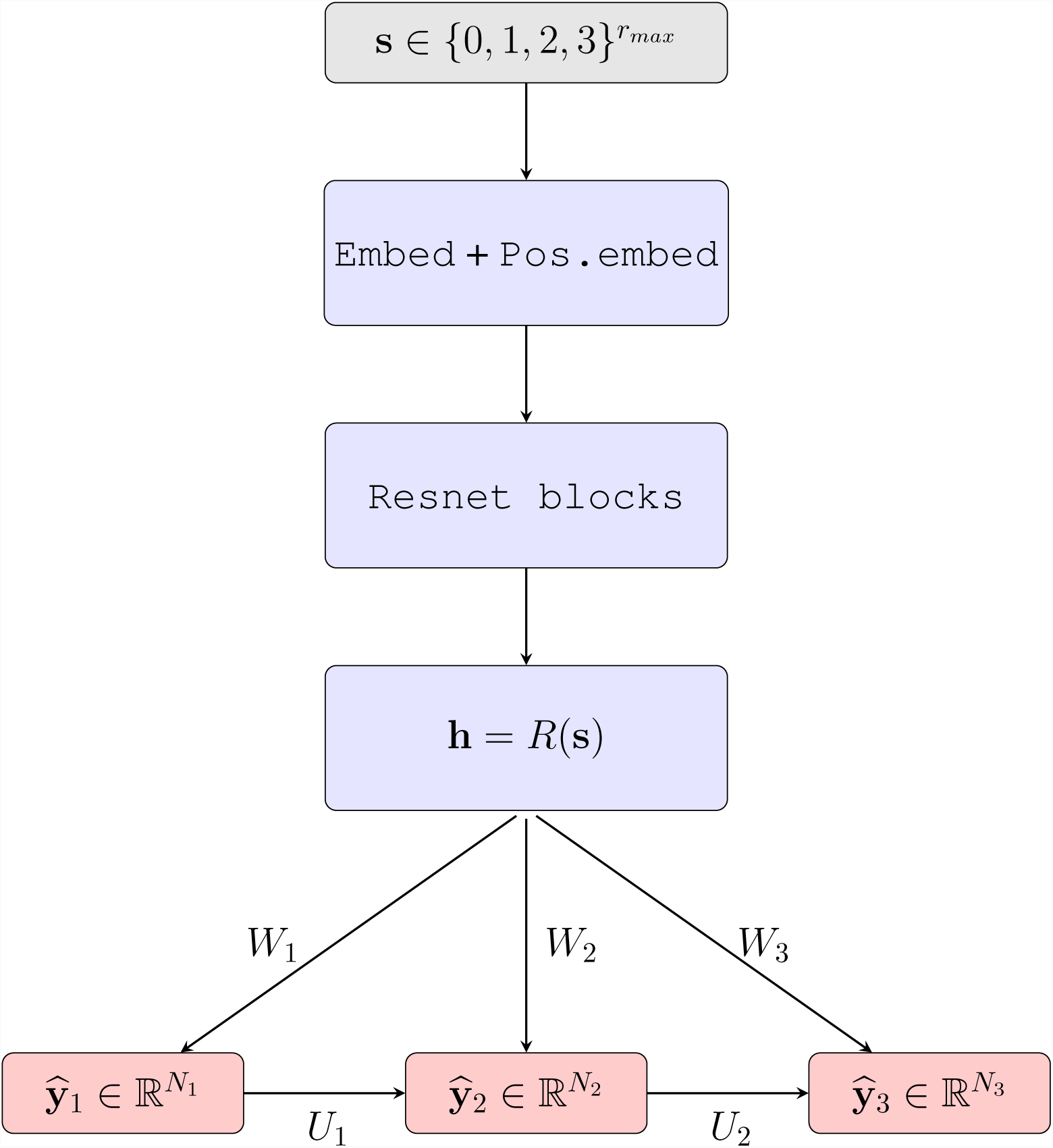
GeNet ised probability architecture. The hidden representation **h** is mapped to *L* softmax layers, representing the unnormal vector over taxa at *L* levels in a hierarchical taxonomy. Here, only 3 levels of the taxonomy are depicted, GeNet is trained with *L* = 7 taxonomic levels.

We perform grid search for some of the hyper-parameters of the network, the ranges considered are in Table 8. We selected the final version of GeNet based purely on validation accuracy on data drawn from the same distribution as the training data.

**Table 8:** Hyperparameters for GeNet.

| Parameter | Values |
| --- | --- |
| Number of initial filters $n_f$ | {128, 256} |
| Size of 1d kernel $v$ | {2, 3, 5} |
| Learning rate $l_r$ | {0.5, 1, 10, 20} |
| Size of fully connected layer $f_c$ | {512, 1024} |
| Batch size $M$ | 64 |

The model used for the experiments, which led to the highest validation accuracy, has the following parameters:

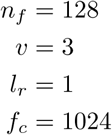

### B. Details for biological software

#### B.1. Generation of Illumina and Pacbio reads

Illumina reads were generated for the *K* genomes used during training with the `RandomReads` module of BBMap, downloaded from https://sourceforge.net/projects/bbmap.

For Illumina reads, we considered four noise parameters: *q ∈*{0.01, 0.1, 0.2, 0.3}. We compute the corresponding Phred quality scores *n* = −10 log_10_(*q*) which are given as input to the `RandomReads` module.

For each noise value, we generate 10 datasets with different random seeds. Each dataset contains *nc* = *N/K* reads of length 1, 000 from each of the *K* genomes observed during training, for *N* = 1, 000, 000. A typical call of the module would look as follows:

~~~
randomreads.sh ref=NC 017449.1.fasta
out=NC 017449.1.fastq len=1000
metagenome=f addpairnum=t reads=nc q=q
seed=dataset num
~~~

For the pathogen experiment in Section 4.2, we generate one dataset with PacBio reads from 21 organisms. A typical command would look as follows:

~~~
randomreads.sh ref=NC 017449.1.fasta
out=NC 017449.1.fastq minlength=1000
maxlength=10000 pacbio=t pbmin=q
pbmax=q metagenome=f addpairnum=t
reads=nc seed=-1
~~~

#### B.2. Obtain ground truth labels with minimap2

We use minimap2(https://github.com/lh3/minimap2) to align the reads from the four Nanopore datasets in Nicholls et al. (2018) to the reference genomes, provided by assembling the reads in these communities using Illumina technologies. The file `Zymo-Isolates-SPAdes-Illumina.fasta.gz` is also provided by Nicholls et al. (2018). Given a FASTQ file with reads `reads.fastq`, we used minimap2 with the following paramters:

~~~
minimap2 -ax -map-ont -t 32
Zymo-Isolates-SPAdes-Illumina.fasta.gz
reads.fastq > align.sam
~~~

#### B.3. Computation of Average Nucleotide Identity (ANI) and Average Amino-Acid Identity (AAI)

We compute ANI using FastANI (https://github.com/ParBLiSS/FastANI) and AAI using CompareM (https://github.com/dparks1134/CompareM).

Given a file `genomes.txt` containing a list of paths to the *K* genomes is the dataset, we ran the following command:

~~~
fastANI -rl genomes.txt -ql genomes.txt -t 8 -o similarity.out.
~~~

This returns a value of zero for many pairs of genomes, since fastANI return zero for values under 70. We completed the similarity matrix using AAI, computed with compareM as follows:

~~~
comparem aai wf --cpus 32 genomes.txt aai.
~~~

## Footnotes

1 We use the terms organism and genome interchangeably for the rest of the paper.

2 Further work could try to split a longer read in smaller pieces, classify each shorter read, and aggregate the results into a classification decision.

3 For Centrifuge, we use the “Bacteria, Archaea, Viruses, Human” database available in https://ccb.jhu.edu/software/centrifuge/. For Kraken, we use the full standard database. We write Kraken instead of Kraken2.

